# *In-vitro* evaluation of Ozenoxacin and other Antibiotics against *Staphylococcus aureus* and *Streptococcus pyogenes* isolated from Skin and Soft Tissue Infections

**DOI:** 10.64898/2026.06.09.731036

**Authors:** Shyam Tripathi, Smita Singhal, Rivan Berlia, Manoj Kumar Yadav, Bansidhar Tarai, Anjali Priyadarshini, Renu Choudhary, Poornima Sheba Samuel Raj, V. Samuel Raj

**Affiliations:** Centre for Drug Design Discovery and Development (C4D), SRM University, Delhi-NCR, Sonepat - 131 029, Haryana, India; Department of Biotechnology and Microbiology, SRM University, Delhi-NCR, Sonepat - 131 029, Haryana, India; Department of Microbiology & Molecular Biology, Max Super Speciality Hospital, Saket, New Delhi 110017, India; Faculty of Management and Commerce, SRM University, Delhi-NCR, Sonepat - 131 029, Haryana, India; PR Institute of Medical Science& Research (PRIMSR), SRM University, Delhi-NCR, Sonepat -131 029, Haryana, India

**Keywords:** antibacterial activity, clinical isolates, Gram-positive, MSSA, MRSA, ozenoxacin, skin and soft tissue infections

## Abstract

*Staphylococcus aureus* and *Streptococcus pyogenes* are major causative bacteria responsible for skin and soft-tissue infections (SSTIs) such as impetigo. Increasing resistance to commonly used topical antibiotics necessitates evaluation of newer agents for the treatment of skin infections. Ozenoxacin, a novel non-fluorinated topical quinolone, has shown promise, exhibiting potent activity against a wide range of pathogens, including methicillin-resistant Staphylococcus (MRSA) and *Streptococcus pyogenes*. The present study compared the *in vitro* activity of ozenoxacin and comparator agents against clinical isolates of *Staphylococcus aureus* and *Streptococcus pyogenes* from multiple sources including skin and soft-tissue, wound, abscess, and blood. Ozenoxacin was assessed for *in vitro* antimicrobial activity against 109 methicillin-susceptible (MSSA), methicillin-resistant *S. aureus* (MRSA), and 24 *Streptococcus pyogenes* isolates by broth microdilution method recommended by the Clinical and Laboratory Standards Institute (CLSI). Ozenoxacin demonstrated potent *in-vitro* activity against all 109 *S. aureus* (MIC_50_/_90_= 0.125/0.5 µg/ml) and 24 *S. pyogenes* (MIC_50_/_90_= 0.015/0.03 µg/ml) strains. In contrast, higher MICs were observed for fusidic acid and mupirocin among a subset of *S. aureus* isolates. A comparison of MIC_90_values demonstrated that ozenoxacin (0.5 µg/ml) was more active against *S. aureus* isolates than 8 of the 9 comparator agents tested including vancomycin and linezolid (MIC_90_= 2 & 4 µg/ml) respectively. *In vitro* studies of ozenoxacin showed potency against staphylococci and streptococci including resistant *S. aureus* strains. These findings support its role as an effective first-in-class quinolone topical therapeutic option in the management of various SSTIs.

## INTRODUCTION

The principal barrier against microbial invasion is the skin. It constantly interacts with the external environment and is colonized with a diverse population of microbes. Most of the colonizing flora consists of bacteria. To help organize the distribution of flora, one can divide the body into two halves at the waistline. The typical organisms that colonize the skin above the waist are usually Gram-positive species such as *Staphylococcus epidermidis, Staphylococcus aureus (S aureus)*, and *Streptococcus pyogenes* and Corynebacterium species. *S. aureus* and *S. pyogenes* species are particularly significant because they contribute to a majority of skin and soft tissue infections (SSTIs) (1). *Staphylococcus aureus (S. aureus*) is both a commensal bacterium that is persistently or transiently present in up to 80% of healthy individuals and a Human pathogen causing a wide spectrum of clinical infections in humans, ranging from superficial skin and soft-tissue infections (SSTIs) to deeply invasive life-threatening bacteraemia and endocarditis.

Skin and soft tissue infections are clinical entities of variable presentation, etiology and severity that involve a broad range of conditions involving the dermis and subcutaneous tissue, extending from minor superficial processes/ infections such as pyoderma to deeper tissue involvement such as necrotizing fasciitis with potentially fatal complications that demand immediate attention (2).

Impetigo is a highly contagious bacterial superficial skin infection, that primarily affects children between 2 to 5 years of age, however, individuals of any age can be susceptible to this bacterial condition (3). It is classified under the broader category of SSTIs which represent a significant proportion of community-and hospital-acquired infections worldwide. Impetigo though common, is often self-limiting, in which systemic symptoms are rare and is an underdiagnosed important paediatric dermatological health problem all over the world, that, if left untreated, can have serious consequences (4).

### Evolving Microbiology and Clinical Implications

The most common pathogens implicated in impetigo are *S. aureus* (including methicillin-resistant *S. aureus* (MRSA) and *S. pyogenes*. Impetigo is a bacterial infection where the causative organism is not routinely identified, however an epidemiological shift from *S. pyogenes* to *S. aureus* has been observed (5). Infection with *S. aureus* or *S. pyogenes* causes the nonbullous type of impetigo which occurs in around 70% of cases, whereas *S. aureus* exclusively causes bullous impetigo, with the production of exfoliative toxins (6). This epidemiologic shift has important clinical consequences since *S. aureus* carries a large repertoire of virulence determinants in conjunction with different antibiotic resistance phenotypes more than *S. pyogenes*. Previously restricted to the hospital environment, *S. aureus* is now more widespread outside healthcare facilities and SSTIs caused by community-associated MRSA (CA-MRSA) have been increasing over the last decade and a serious problem worldwide (7).

Topically applied antibiotics, such as mupirocin and fusidic acid, have been the mainstay/first-line of treatment for superficial SSTIs and localized impetigo, although retapamulin is a more recent alternative (8). The development of new antibiotics represents a current need, considering the increase in multidrug-resistant pathogens, such as MRSA and mupirocin-resistant or quinolone-resistant Staph aureus, raising concerns regarding their continued effectiveness (9). Moreover, fusidic acid monotherapy, specifically topical formulations, has been strongly linked to the emergence of fusidic acid resistance among either MRSA or methicillin-susceptible Staph aureus (MSSA) (10). This increasing resistance complicates treatment options and highlights the need for newer topical antimicrobials with improved efficacy and a lower propensity for resistance development (11).

The most recent topical option ozenoxacin a novel nonfluorinated quinolone with bactericidal activity, is approved in the USA, Canada and 12 EU countries as a 1% cream formulation for treatment of impetigo in children and adults and in Japan as a 2% lotion formulation for treatment of superficial skin infections and acne. Approval of ozenoxacin was granted based on demonstrable clinical benefit in two large Phase III trials of impetigo involving adults and children from 2 months of age (8). Rapid eradication of bacterial load is important in impetigo to hasten symptom resolution and limit person-to-person transmission of infection.

Usually, gyrase inhibitors are more active against DNA gyrase than topoisomerase, while quinolones show more inhibitory potency against Topoisomerase IV than that of Gyr B (12). Interestingly, ozenoxacin’s dual inhibitory activity against DNA gyrase (topoisomerase II), and topoisomerase IV, required for bacterial DNA replication will pose lower risks of resistance development than single-target inhibitors (13). Lack of a fluorine atom at the C-6 position theoretically has less phototoxicity potential than traditional fluoroquinolones which contributes to its favourable safety profile and reduced systemic absorption, and is therefore considered suitable for paediatric use.

Ozenoxacin has potent antibacterial activity against *S. aureus* (MSSA, MRSA) and *S. pyogenes* irrespective of their levofloxacin susceptibility status. The present study aimed to evaluate the *in-vitro* antibacterial activity of ozenoxacin against clinical isolates of *S. aureus* (including MRSA and MSSA) and *S. pyogenes* obtained from SSTIs and to compare its activity with commonly used topical agents and selected systemic antibiotics.

### Study Rationale and Objectives

In view of the increasing incidence of resistant pathogens, *in vitro* assessment of the activity of ozenoxacin against recent clinical isolates is important. This study aimed to:

1. Assessment of the *in vitro* antibacterial activity of ozenoxacin against clinical isolates of *S. aureus* (including MRSA) and *S. pyogenes* obtained from patients with skin infections and other sources.
2. Compare the efficacy of ozenoxacin with that of other commonly used topical antimicrobial agents, and relevant systemic antibiotics.
3. To assess the susceptibility patterns of these clinical isolates, including the presence of resistance to existing topical treatments.

## MATERIALS AND METHODS

### Study Design and Clinical Isolates Collection

The study was conducted in the Bacteriology lab in Centre for Drug Design Discovery and Development (C4D) at SRM University Delhi NCR, Sonipat, Haryana. Ethical approval was obtained from the Institutional Ethics Committee. This study involved bacterial isolates and the isolates were collected from the microbiology laboratories. This was a laboratory-based, cross-sectional *in-vitro* study designed to evaluate the antibacterial activity of ozenoxacin against Gram-positive clinical isolates associated with skin and soft tissue infections and other sources.

A total of 133 clinical isolates, including *Staphylococcus aureus* (n=109), and *S. pyogenes* (n=24), were included in this study. These isolates include American Type Culture Collection (ATCC; Manassas, VA, USA) strains, and clinical isolates obtained from Indian hospitals were also used in *in vitro* screening. Gram-positive bacteria included in the study as principal pathogens were *Staphylococcus aureus* clinical strains obtained from the culture bank at C4D. The culture bank comprises well-characterized clinical isolates recovered from specimens submitted by tertiary care hospitals and accredited diagnostic laboratories. Isolates were sourced from patients and only one isolate per patient was included to avoid duplication (non-duplicate) in the study. A total of 109 *S. aureus* and 24 *Streptococcus pyogenes* clinical isolates were profiled for their *in-vitro* susceptibility to ozenoxacin and comparator antimicrobial agents.

Each isolate was identified at the referring hospital/laboratory and added to the culture collection in C4D, SRM University, Sonepat. Each isolate was accompanied by relevant data, including genus and species name. Upon receipt, isolates were sub-cultured on to blood agar to ensure viability and purity. Methicillin resistance among *S. aureus* isolates was determined using Cefoxitin susceptibility testing in accordance with CLSI guidelines, and isolates were categorized as methicillin-resistant (MRSA) or methicillin-sensitive (MSSA). Strains were stored at −80°C in 20% glycerol in Trypticase soy broth (TSB) (Becton, Dickinson and Company, Cockeysville, MD).

### Media, reagents, and antimicrobial agents

#### Investigational Product

The investigational product i.e. ozenoxacin IP (Batch No. WS/EGY/22?01) and comparator drugs/ antibiotics fusidic acid (Batch No. L12A424901), mupirocin (Batch No. L324223996), amoxycillin clavulanic acid (ACV-1123228) and clindamycin (P-003-CN20230603) were kindly provided by Walter Bushnell Enterprises Private Limited. Remaining antibiotics i.e. vancomycin, linezolid, levofloxacin, neomycin and cefoxitin were procured from Sigma-Aldrich Co. LLC, USA, and Hi-Media, India. All media and reagents were purchased from Becton Dickinson and Company and HI Media.

#### *In vitro* Antimicrobial Susceptibility Testing

Minimum inhibitory concentration (MIC) of ozenoxacin and other comparator drugs was determined against Gram-positive bacterial strains using broth microdilution method, as recommended by the Clinical and Laboratory Standards Institute (CLSI) guidelines. Susceptibility was determined using breakpoints set by the CLSI, except that British Society for Antimicrobial Chemotherapy (BSAC) breakpoints were used for fusidic acid and mupirocin against staphylococci. Concentrations of antibacterial agent tested for the isolates are shown in Table 1. Quality control was ensured using *S. aureus* ATCC 29213, *S. pneumoniae* ATCC 49619 and *Escherichia coli* ATCC 25922 as standard reference strains recommended by CLSI. All quality control results were within the ranges specified by the CLSI.

**Table 1:** Concentrations of antibacterial agents tested for ***S. aureus* and *S. pyogenes* strains**.

| Antibacterial agents | Antibacterial range tested (µg/ml) |  |
| --- | --- | --- |
|  | *Staphylococcus aureus* | *Streptococcus pyogenes* |
| **Ozenoxacin** | 0.004–2 | 0.004–2 |
| **Fusidic Acid** | 0.03–16/ 0.125–128 | 0.03–16 |
| **Mupirocin** | 0.03–16/ 0.125–128 | 0.03–16 |
| **Neomycin** | 0.006–32 | 0.03–16 |
| **Amoxycillin clavulanic acid** | 0.125–64 | 0.125–64 |
| **Clindamycin** | 0.06–32 | 0.03–16 |
| **Linezolid** | 0.03–16 | 0.03–16 |
| **Levofloxacin** | 0.006–32 | 0.03–16 |
| **Vancomycin** | 0.03–16 | 0.03–16 |
| **Cefoxitin** | 0.125–64 | Not tested |

## Statistical Analysis

To formally assess whether ozenoxacin has lower anti-microbial activity thresholds than the comparator drugs, the MIC values for each antibiotic were first log_2_-transformed to put them on a roughly linear scale.

For each antibiotic we calculated the median log_2_(MIC) and interquartile range (IQR). Pairwise two-sided Mann–Whitney U tests were performed in IBM SPSS Statistics (v.25) to compare the distribution of MIC values for ozenoxacin versus each other antibiotic. Effect sizes were calculated as r = Z/√N, where Z is the standardized test statistic and N is the total sample size. To control the familywise error rate across the nine tests, p-values were Bonferroni-adjusted. A Bonferroni-adjusted p < 0.0056 was considered statistically significant.

## RESULTS

### Comparative Antimicrobial activity against Gram-positive bacteria

Ozenoxacin is a novel non-fluorinated quinolone that is being developed for topical use. In this study the *in vitro* activity of ozenoxacin was evaluated against Gram-positive isolates primarily *S. aureus*. In addition, ozenoxacin was also evaluated against *Streptococci pyogenes* clinical strains to see the trend and expand with more strains in future studies. The *in vitro* activities of ozenoxacin against Gram-positive bacteria such as *S. aureus* and *S. pyogenes* are summarized in Table 2

### *In vitro* Activity of Ozenoxacin against Gram-positive Pathogens

#### Staphylococci aureus

Ozenoxacin exhibited good *in vitro* activity against all 109 *S. aureus* strains (MIC_50_/_90_ = 0.125/0.5 µg/ml) respectively (Table 2). Ozenoxacin retained activity against *S. aureus* including MRSA, regardless of methicillin resistance or levofloxacin nonsusceptibility, indicating that its antibacterial activity was largely unaffected by these resistance mechanisms.

**Table 2:** Minimum inhibitory concentration (MIC) data of Ozenoxacin stratified by Methicillin resistance / susceptibility and/or Levofloxacin susceptibility/nonsusceptibility for Gram-positive bacteria (*Staphylococcus aureus*and *Streptococcus pyogenes)*

| Organisms | Levofloxacin susceptibility | Number of strains (n) | Ozenoxacin MIC ($\mu\text{g/ml}$) | | |
| --- | --- | --- | --- | --- | --- |
|  |  |  | MIC<sub>50</sub> | MIC<sub>90</sub> | Range |
| *Staphylococci aureus* | All | 109 | 0.125 | 0.5 | 0.004–2 |
|  | Levofloxacin non susceptible | 95 | 0.125 | 0.5 | 0.015–2 |
|  | Levofloxacin susceptible | 14 | 0.006 | 0.125 | 0.004–0.25 |
| Methicillin-resistant *S. aureus* (MRSA) | All | 85 | 0.125 | 0.5 | 0.004–2 |
|  | Levofloxacin non susceptible | 80 | 0.125 | 0.5 | 0.015–2 |
|  | Levofloxacin susceptible | 5 | 0.004 | 0.015 | 0.004–0.015 |
| Methicillin-susceptible *S. aureus* (MSSA) | All | 24 | 0.125 | 0.25 | 0.004–0.5 |
|  | Levofloxacin non susceptible | 15 | 0.125 | 0.25 | 0.03–0.05 |
|  | **Levofloxacin susceptible** | **9** | 0.008 | 0.015 | 0.004–0.25 |
| *Streptococcus pyogenes* | **All** | **24** | 0.015 | 0.03 | 0.004–0.125 |
|  | **Levofloxacin non susceptible** | **1** | 0.125 | 0.125 | 0.125 |
|  | **Levofloxacin susceptible** | **23** | 0.015 | 0.03 | 0.004–0.03 |

A comparison of MIC_90_values demonstrated that ozenoxacin (0.5 µg/ml) was more active against *S. aureus*isolates than 8 of the 9 comparator agents tested including vancomycin and linezolid (MIC_90_=2 & 4 µg/ml); fusidic acid (MIC_90_=8 µg/ml); levofloxacin, neomycin, amoxicillin-clavulanate and clindamycin (64 µg /ml); and cefoxitin 128 µg/ml). Mupirocin was the only comparator tested that had a MIC_90_value (0.5 µg/ml) equivalent to that of ozenoxacin (Table 3).

**Table 3:** Comparative minimum inhibitory concentration data of Ozenoxacin and other antibacterial agents for Gram-positive isolates from skin and soft tissue infections.

| Organisms<br>(no of strains) | MIC<br>µg/ml | Antimicrobial agent(s) |  |  |  |  |  |  |  |  |  |
| --- | --- | --- | --- | --- | --- | --- | --- | --- | --- | --- | --- |
|  |  | OZN | FUS | MUP | VAN | LZD | LVX | NEO | AMC | CLI | CFX |
| *Staphylococcus aureus* (n=109) | MIC<sub>50</sub> | **0.125** | 0.25 | 0.25 | 1 | 2 | 8 | 4 | 16 | 0.125 | 32 |
|  | MIC<sub>90</sub> | **0.5** | 8 | 0.5 | 2 | 4 | 64 | 64 | 64 | 64 | 128 |
| *Streptococcus pyogenes* (n = 24) | MIC<sub>50</sub> | **0.015** | 8 | 0.125 | 0.5 | 1 | 0.5 | 16 | 0.03 | 0.03 | ND |
|  | MIC<sub>90</sub> | **0.03** | 8 | 0.25 | 0.5 | 1 | 0.5 | >16 | 0.125 | 4 |  |
| AMC: Amoxicillin/Clavulanate; FUS: Fusidic acid; LVX: Levofloxacin; LZD: Linezolid; MUP: Mupirocin; NEO: Neomycin; OZN: Ozenoxacin; VAN: Vancomycin. MIC: Minimum inhibitory concentration; ND: Not done |  |  |  |  |  |  |  |  |  |  |  |

### Activity of Ozenoxacin against *Streptococcus pyogenes*

Ozenoxacin exhibited low MIC values (MIC_50_/_90_=0.015/0.03 µg/ml) against *S. pyogenes*, indicating high potency. Ozenoxacin was highly active against levofloxacin non susceptible/susceptible *S. pyogenes* with a MIC_90_of 0.03µg/ml (Table2). Against *S. pyogenes*, ozenoxacin had the lowest MIC_90_= 0.03 µg/ml of all other antibacterial agents tested. Whereas mupirocin showed two dilutions higher MIC _90_0.25 µg/ml and another topical agent fuscidic acid had much higher MIC_90_8 µg /ml in comparison to ozenoxacin. Neomycin had least activity against *S. pyogenes* MIC_90_>16 µg /ml. MIC _90_values for other comparator agents were higher than that of ozenoxacin: 0.125 µg /ml for amoxicillin-clavulanic acid: 0.5 µg /ml for vancomycin and levofloxacin and 1µg/ml for linezolid (Table 3).

## Statistical Analysis

The results also indicate that the mean MIC of ozenoxacin is lower than of all other antibiotics considered for analysis. The median log_2_(MIC) for ozenoxacin was –3.00 (IQR –4.06 to –2.00) across all isolates. Comparatively, the other antibiotics had higher median log_2_(MIC) values, indicating lower potency (higher MIC): e.g. fusidic acid –2.00 (–3.00 to –1.00), mupirocin –2.00 (–2.00 to – 1.00), vancomycin 0.00 (–1.00 to 0.00), levofloxacin 3.00 (2.00 to 5.00), etc. Pairwise Mann–Whitney tests showed that ozenoxacin MIC values were significantly lower than those of all other studied antibiotics except Clindamycin. For example, against cefoxitin the U statistic was 2.0 (Z = –12.753, p<0.001), and the rank-biserial effect size was r = 0.86. Similarly, the comparisons versus linezolid (U=236.0, Z = –12.251, p<0.001, r=0.83), amoxicillin–clavulanate (U=234.0, Z = –12.255, p<0.001, r=0.83), and vancomycin (U=1037.5, Z = –10.529, p<0.001, r=0.71) all yielded highly significant differences (Bonferroni-corrected p<0.001). In contrast, clindamycin had an almost identical median MIC to ozenoxacin (both –3.00) and the test was non-significant, indicating no appreciable difference in potency (Table 4).

**Table 4:** Statistical output using Pairwise Mann–Whitney U tests comparing Ozenoxacin vs. each antibiotic (n=109 each group) against *S. aureus*.

| Comparator Antibiotics | Mann-Whitney U test | Z | r | p-value* |
| --- | --- | --- | --- | --- |
| **Fusidic acid** | 3850.5 | $-4.488$ | 0.3 | **< 0.001** |
| **Mupirocin** | 3266.5 | $-5.743$ | 0.39 | **< 0.001** |
| **Vancomycin** | 1037.5 | $-10.529$ | 0.71 | **< 0.001** |
| **Linezolid** | 236 | $-12.251$ | 0.83 | **< 0.001** |
| **Levofloxacin** | 506.5 | $-11.670$ | 0.79 | **< 0.001** |
| **Neomycin** | 570.5 | $-11.532$ | 0.78 | **< 0.001** |
| **Amox clavulanate** | 234 | $-12.255$ | 0.83 | **< 0.001** |
| **Clindamycin** | 5265.5 | $-1.450$ | 0.1 | Not significant |
| **Cefoxitin** | 2 | $-12.753$ | 0.86 | **< 0.001** |
| r is the effect size ($Z/\sqrt{N}$). p-values are two-sided; $p_{\text{adj}}$ = Bonferroni-corrected.<br>*Significant (Bonferroni-adjusted) p-values are in bold. | | | | |

Because Mann–Whitney tests rank-based differences rather than assuming normality, these results are robust to outliers and non-normal MIC distributions. The use of Bonferroni correction ensured a conservative control of type I error across multiple tests. Even under this strict criterion, ozenoxacin was significantly more potent than virtually all comparators. These findings support the interpretation that ozenoxacin exhibits significantly stronger antimicrobial activity (lower MIC) relative to the other antibiotics in this study.

## DISCUSSION

Ozenoxacin, a novel nonfluorinated quinolone with bactericidal activity, is approved in the USA, Canada and 12 EU countries as a 1% cream formulation for treatment of impetigo and in Japan as a 2% lotion formulation for treatment of superficial skin infections and acne. During development of ozenoxacin, surveillance studies compared it’s *in vitro* antibacterial activity with that of other topical and systemic antibacterial agents (8). The current comparative study examined the antibacterial activity of ozenoxacin in Gram-positive clinical isolates derived mainly from SSTIs, RTIs and UTIs, which were stratified according to their methicillin and levofloxacin susceptibility status. The antibacterial activity of ozenoxacin was compared with that of 9 other antibacterial agents against 133 Gram-positive isolates.

The results of this study demonstrate that ozenoxacin possesses potent *in-vitro* antibacterial activity against Gram-positive clinical isolates from SSTIs, including MRSA and *S. pyogenes*. This is in line with results of previously published comparative studies wherein ozenoxacin was shown to be a potent antibacterial agent against Gram-positive (Staphylococci and Streptococci) isolates (8).

The low MIC values observed indicate superior activity compared to commonly used topical antibiotics such as mupirocin and fusidic acid. Statistical significance results are observed with p-value of (<.0001) for all comparator antibiotics except clindamycin against the MSSA and MRSA clinical isolates. Resistance to mupirocin and fusidic acid has been increasingly reported, particularly among *S. aureus* isolates. In this study, elevated MICs for these agents were observed in a subset of isolates, highlighting the clinical relevance of alternative topical therapies.

Ozenoxacin’s dual inhibition of DNA gyrase and topoisomerase IV, along with its non-fluorinated structure, may contribute to its enhanced activity and reduced potential for resistance development. The observed efficacy against MRSA is particularly significant, given the limited topical options available for such infections (14). Although linezolid and levofloxacin demonstrated good *in-vitro* activity, their systemic use and potential adverse effects limit their applicability as first-line agents for superficial SSTIs. Therefore, there is a high unmet medical need for a new class of anti-MRSA antibiotic to overcome the limitations of SOC and fight against emerging resistance (12).

## LIMITATIONS

This study was limited to *in-vitro* analysis and did not assess clinical outcomes. Additionally, pharmacokinetic and pharmacodynamic correlations were not evaluated. Further clinical studies are warranted to corroborate these findings.

## CONCLUSION

In this study, the *in vitro* activity of ozenoxacin against Gram-positive clinical isolates is compared with a panel of commonly used antibacterial agents and ozenoxacin was shown to be a potent antibacterial agent against *S. aureus* and *S. pyogenes* isolates. The excellent activity of ozenoxacin in staphylococcal and streptococcal spp. including methicillin and quinolone-resistant strains most commonly associated with childhood impetigo suggests that ozenoxacin may be a valuable alternative to other topical antibiotics in the topical management of SSTIs and underscores its importance in antimicrobial stewardship initiatives.

## CONFLICT OF INTEREST

All the authors declare that there is no conflict of interest.

## AUTHORS’ CONTRIBUTION

All authors listed have made a substantial, direct, indirect, and intellectual contribution to the work, and approved it for publication.

All authors have read and agreed to the published version of the manuscript.

## FUNDING

This study was financially supported by Walter Bushnell Enterprises Pvt Ltd. Gurgaon, India through a service agreement. The study also received material support in the form of study drugs and antibiotics.

